# High-frequency common inputs entrain motoneuron subpopulations differently

**DOI:** 10.64898/2026.03.24.713960

**Authors:** Alejandro Pascual Valdunciel, Javier Yanguas Mayo, Emanuele Abbagnano, Natalia Torres Cónsul, Filipe Nascimento, M. Görkem Özyurt, Dario Farina, Jaime Ibáñez

**Author notes:** Equal Contribution (Dario Farina, Jaime Ibáñez).

## Abstract

Spinal motoneuron (MN) pools behave as linear systems that transmit common synaptic input to muscles. However, MNs are biophysically heterogeneous and intrinsically nonlinear. How different MN subpopulations integrate and transmit high-frequency inputs remains poorly understood, partly because conventional analyses treat the MN pool as a single functional system rather than examining subpools with different firing rates. Here, we addressed this gap using a combination of computational simulations and human MN recordings. Simulations of MNs receiving a common synaptic input at varying frequencies showed that MNs’ firings become phase-locked to input oscillations when the input frequency approximates the neuron’s firing rate or its harmonics. We refer to this frequency-dependent synchronization as entrainment. Importantly, this subpool-specific effect was masked when MN activity was analysed at the whole-pool level. Because entrained MNs effectively sample the input at their firing instants, we developed a MN-firing locked method that uses individual MN firings as endogenous triggering events for peristimulus frequencygrams across the pool. In simulations, this method revealed entrainment-driven firing rate modulations across MN subpools. We then applied this MN-firing locked method to MNs decomposed from high-density surface electromyography recordings obtained during isometric contractions in healthy individuals. We found that faster-firing MNs exhibited larger transient firing rate increases, time-locked to slower MN activity. Furthermore, these modulations correlated with common input in the alpha and beta bands implicating high frequency common input as the driving source. Together, these findings demonstrate that MN nonlinearities generate heterogeneous, frequency-dependent dynamics that remain hidden in conventional pool-level analyses.

## INTRODUCTION

Spinal α-motoneurons (MNs) are specialized neurons located in the spinal cord that integrate synaptic inputs from the central and peripheral nervous systems and project peripherally to innervate skeletal muscle fibers, thereby coordinating muscle activity (Heckman *et al*., 2009). Identifying the sources and quantifying the synaptic strength of these inputs is essential for understanding how the nervous system organizes muscle activity, particularly during tasks requiring precise motor control or sustained force generation (Myers *et al*., 2004).

MNs exhibit heterogeneous intrinsic properties and operate as non-linear systems, in which intrinsic features such as membrane conductance determine how synaptic input is converted into action potential trains at specific firing rates, which are then transmitted to muscle fibres. Despite this individual-level non-linearity, when considered collectively, the MN pool behaves as a linear amplification system of the input commonly projected to all MNs (Farina & Negro, 2015; Watanabe & Kohn, 2015). This linear-transformation of common input applies to both low-frequency components (<10 Hz), which represent the effective neural drive to the muscles transformed into force, as well as to higher input frequencies, such as alpha (8-12 Hz) or beta (13-35 Hz) bands, which have limited or negligible effect on force modulation (Zicher *et al*., 2023). Despite their limited contribution to force control, alpha and beta band inputs are well-characterized components of descending corticospinal drive during voluntary contractions (Farmer *et al*., 1993; Baker, 2007) and are implicated in motor disorders such as Parkinson’s disease and essential tremor (Benito-León *et al*., 2019; Vinding *et al*., 2020).

Most studies characterizing common input exploit this pool-level linearization by summing the activity of a sufficiently large number of MNs combined with the application of spectral methods such as coherence analysis (Negro & Farina, 2012*a*). However, because MNs process inputs differently due to their distinct intrinsic properties, pooling MN activity inevitably discards information about the behaviour of functionally distinct subpopulations (Farina *et al*., 2014). This loss of information is relevant due to different reasons. First, motor unit decomposition often yields partial populations with different types of motor units that have variable firing characteristics (Caillet *et al*., 2023). Second, selective affectation of specific MN types (e.g., fast vs slow-type MNs) may not be captured if the focus of a study relies only on the output of the whole motor pool, rather than individual subpool elements (Huh et al., 2021). It is therefore important to understand how the properties of individual MNs and subpools of MNs affect the characterization of the neural inputs projected to muscles.

To investigate the characteristics of individual MN responses to common input, it is necessary to move beyond the MN pooling approach typically used in coherence analysis. Time-domain methods such as peristimulus frequencygram (PSF) can be utilized to characterize the timing and amplitude of synaptic inputs arriving at individual MNs (Türker & Cheng, 1994). Classical PSF methods rely on delivering external stimuli to synchronize MN activity with controlled evoked common input. While powerful for studying reflex pathways, these methods cannot characterize the transformation of endogenously generated common inputs during voluntary contractions, as they rely on precise temporal events that are unknown in the absence of external stimuli (Yavuz *et al*., 2018; Pascual-Valdunciel *et al*., 2025). Thus, new analytical approaches are needed to characterize the timing and amplitude of common inputs in more naturalistic conditions, that are sensitive enough to account for individual MN properties across motor pools.

Computational studies have shown that when the frequency of an oscillatory common input approximates a MN’s firing rate, the MN’s discharges tend to align with the peaks of the excitatory input (Hutcheon & Yarom, 2000; Lowery & Erim, 2005). This synchronization of MU firing activity with the common input oscillations — here termed entrainment — may be different across MN subpools, due to MN heterogeneous intrinsic properties that are thus reflected in different firing rates. If firing rates determine how individual MNs sample common inputs, then the spiking activity of MNs will reveal information about that common input. Because MNs that are entrained (due to a relationship between the input and their firing rate) will tend to fire aligned with the common input, their spiking activity can serve as endogenous triggering events for PSF analysis. When applied across large populations of MNs with different firing rates, this MN-firing locked approach would reveal how common inputs at various frequencies are sampled differently by distinct MN subpools.

Here, we combined computational MN simulations and human MN recordings to investigate how common inputs produce heterogeneous firing dynamics across MN populations. We first use computational simulations to establish that MN subpopulations become differently entrained to common oscillatory input as a function of their firing rate, and that this effect is masked when MN activity is analysed collectively. Building on this, we develop a MN-firing locked method in which individual MN firing times serve as endogenous triggers for PSF analysis and validate this method’s ability to detect differential firing rate modulations across MN subpools driven by the common input. Finally, we apply the MN-firing locked method to experimental high-density surface electromyography (HDsEMG) data from healthy subjects, revealing that high-frequency common inputs produce firing rate dynamics that differ systematically across MN subpools—a heterogeneity that population-level methods cannot detect. Together, these findings provide a framework for extending synaptic input analysis beyond population-level measures, particularly for resolving the behavior of distinct MN subpopulations.

## METHODS

### Motoneuron simulations

We implemented a biophysical model of a MN built upon Hodgkin-Huxley equations with both dendritic and somatic compartments (Cisi & Kohn, 2008). The goal was to study the response of different MN pools to different common input frequencies. MN pools were created by linearly interpolating experimental cellular MN properties from the smallest to largest MN in line with the size principle (Henneman, 1957). The MN parameters were derived from *in vivo* MN recordings obtained from S-type (slow twitch) MNs. A pool of 80 MNs was simulated across different conditions in which MNs received different common excitatory input profiles. Membrane voltage dynamics for each MN were calculated by solving the differential equations described in Equation 1 using a fourth order Runge Kutta integration method with time steps of 0.2 ms. In this equation, *C*_*m*_ denotes membrane capacitance and *I*_*ext*_ represents the external synaptic current delivered to the MN. For each ion channel or synaptic conductance *k*, the parameter *g*_*k*_ specifies its maximal conductance, *v* indicates the membrane voltage of the compartment, and *E*_*k*_ defines the corresponding reversal potential for that conductance. A more detailed description of the ion channel equations and model parameters is provided in (Pascual-Valdunciel *et al*., 2025).

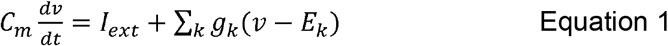

The simulations reproduced MN behavior during sustained firing, similarly to an isotonic and isometric contraction performed by an individual. Then, the input to the MN pool was defined as follows:

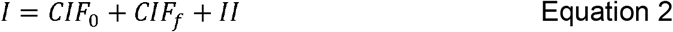

In our simulations, the input (I) was the combination of three inputs: (1) the common input CIF_0_, that refers to the effective neural drive that contributes to stable force generation and produces correlated fluctuations between MNs, as it is uniformly distributed within the pool (Farina & Negro, 2015); (2) the independent noise each MN receives (*II*); and (3) an additional common input delivered at given frequency band (*CIF*_*f*_) (**Figure 1A)**. *CIF*_*0*_ was represented by a constant synaptic current plus a white noise low-passed filtered below 5 Hz with amplitude equivalent to 10% of the total amplitude. Importantly, by doing a hyperparameter sweep, the amplitude of *CIF*_*0*_ was tuned so the firing rate of the resulting MNs would match that of the experimental dataset (approximately 10-16 Hz). *II* was modelled as white noise that follows a distribution of N (0,1 σ^2^) normalized to 30% of the amplitude of *CIF*_*0*_ (Binder & Powers, 2001). *CIF*_*f*_ was modelled as a synaptic current composed of white noise band-pass filtered at a specific frequency *f* with a bandwidth of ± 2 Hz. The frequency *f* varied from 2 to 50 Hz across simulations, with 1 Hz increments, and the total power of the current was kept constant across simulations with different frequencies. An additional condition was simulated with the MN pool devoid of any *CIF*_*f*_ input (only *CIF*_*0*_). All the simulations consisted of an increasing input ramp of 2s until reaching the target common input level, followed by an 80 s plateau, and a finishing with 2 s falling phase.

**Figure 1.**
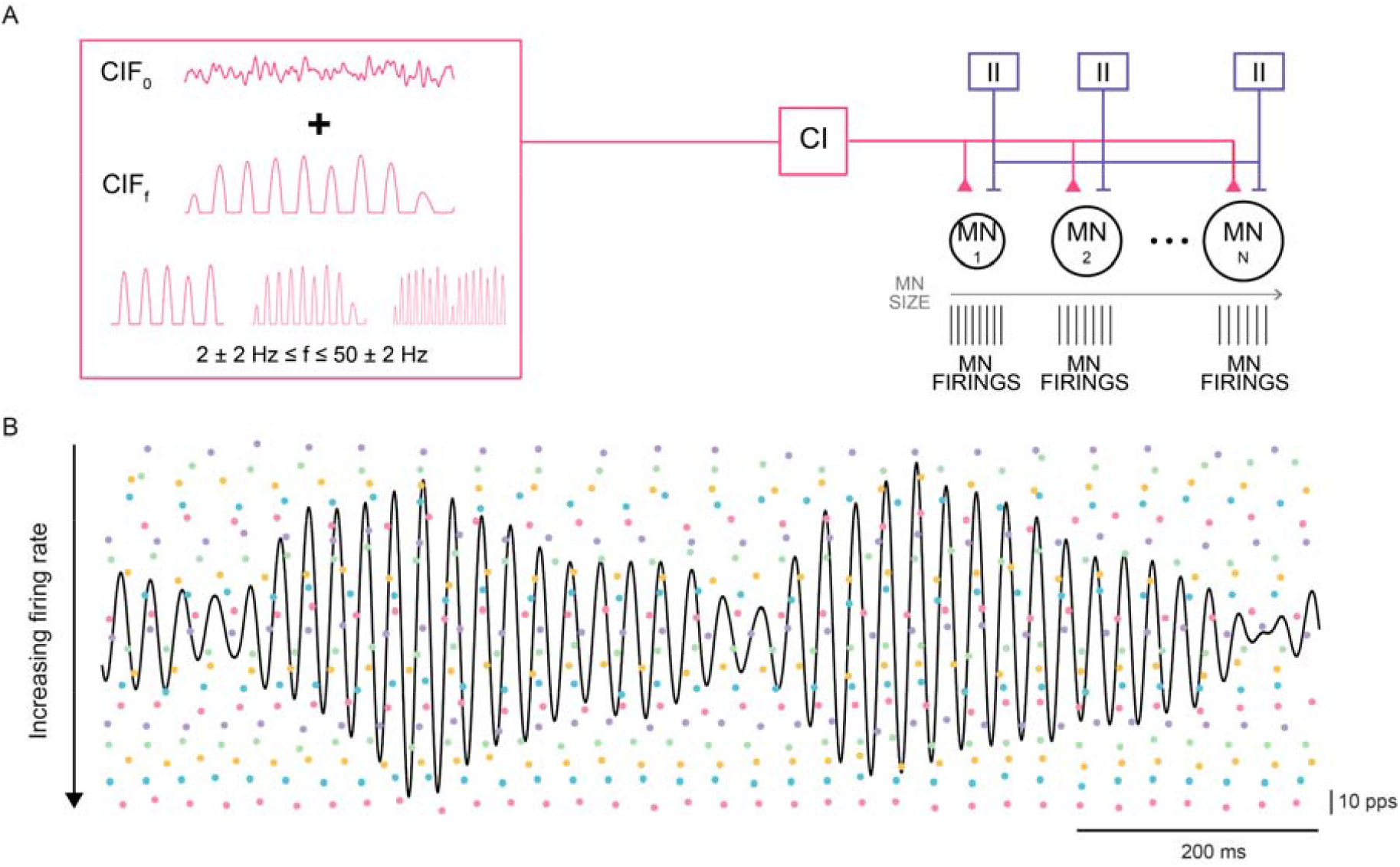
MN simulations. **A**. Schematic of the MN simulations. **B**. Example of MN spike train output from the model. Solid colored dots represent firings of individual MNs with different sizes (increasing size from bottom to top; information on instantaneous firing rates is encoded on the vertical axis for each firing). Black solid line represents the high frequency common input (CIF_f_ delivered at 20 Hz) received by all the MNs.

The spike trains corresponding to 55 s of the plateau of the simulation were selected for analysis. The selection of this window was based on matching the duration of the experimental recordings. Then, the instantaneous firing rate (iFR) and its average 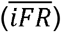 were computed for each MN (Del Vecchio *et al*., 2020). Consequently, MNs were grouped in quartiles based on firing rate (**Figure 1B)**. In this study, we focused on the results of the first and last quartile, representing the active MNs in the pool with the fastest and slowest firing rates.

### MN entrainment and MN-firing locked analysis

We implemented a time-domain method to investigate how common input shapes firing dynamics across MN subpools. Considering that the firing times of individual MNs contain sampling information of the input, this method is based on PSF built by time-locking the firing activity of one MN to the firing events of other individual MNs within the same pool, allowing detection of transient firing rate modulations.

This analysis was applied to all the available pairs of MNs per simulation, and later per subject and contraction level. The instantaneous firing rate (*iFR*) and firing times of individual MNs were used to time-lock the firing events of the rest of the MNs in a pair-based fashion to build PSFs and peristimulus time histograms (PSTH) (**Figure 2A, B**). For a given pool of *N* MNs, the *iFR* and *firing instants* for the *j*^*th*^ MN (*MN*_*j*_) were used to align the *iFR* (for PSF) and firing counts (for PSTH) of all remaining MNs (*MN*_*j+1…N*_), within 200 ms windows centred on each firing event. Then, PSF was built by superimposing (summing for PST) the segmented windows of *iFR* across *M* firings available for the triggering MN (*MN*_*j*_) and binning them in bin widths of 0.5 ms. Afterwards, the cumulative sums (CUSUM) of PSF and PSTH were calculated using equations 3 and 4.

**Figure 2.**
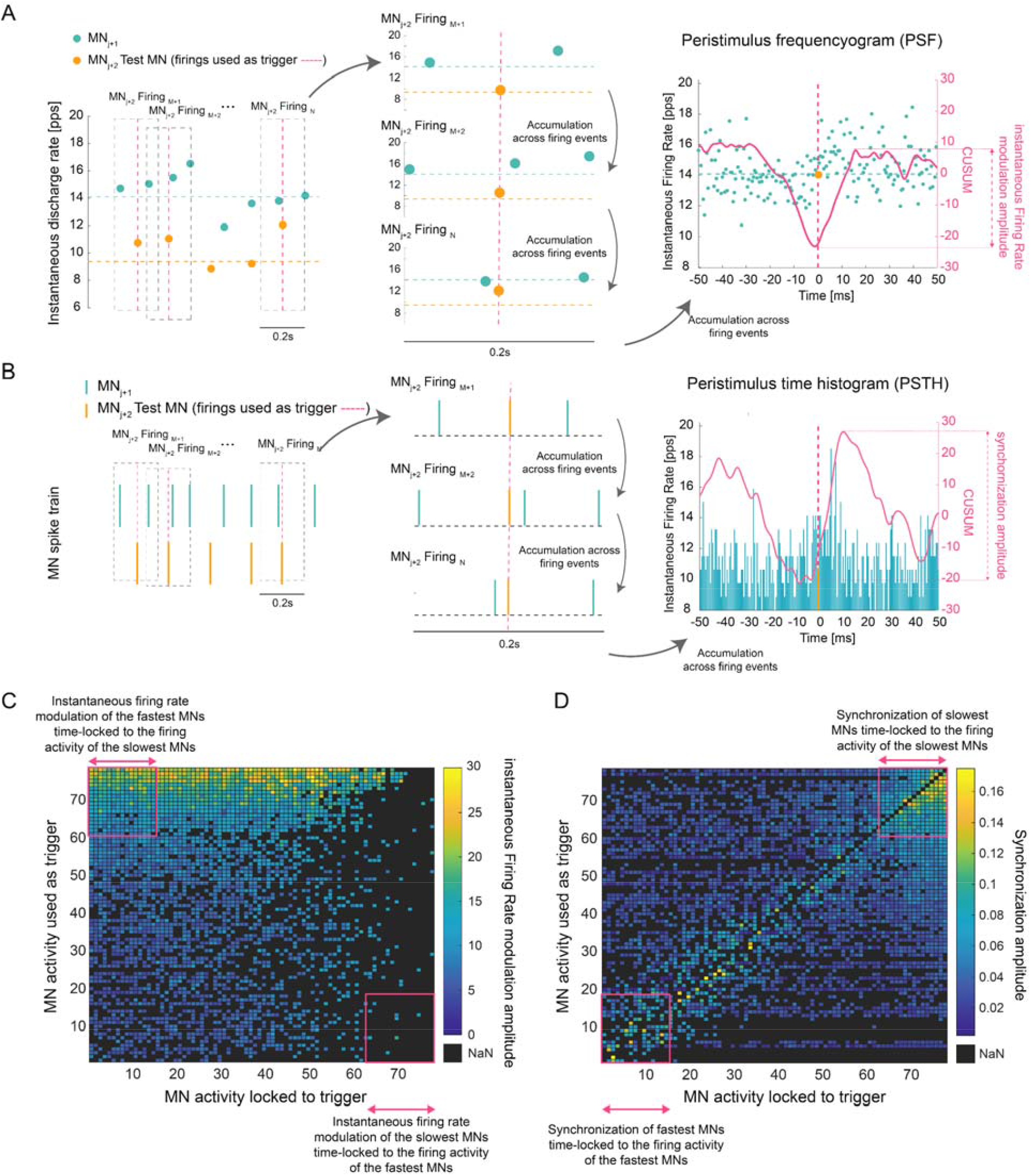
MN-firing locked analysis. **A**. MN-firing locked PSF for a pair of MNs to estimate firing rate modulation. **B**. MN-firing locked PSTH for a pair of MNs to estimate synchronization. **B**. MN-firing locked PSTH for a pair of MNs to estimate MN synchronization. **C, D**. Heatmap of the instantaneous firing rate modulation (**D**) and synchronization amplitude (**C**) across MN pairs (data from one simulation with CIF_f_ delivered at 10 Hz). MNs are sorted by

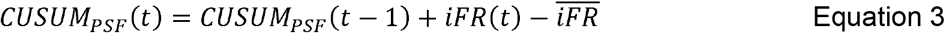

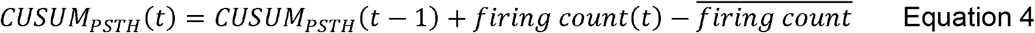

This process was repeated for each MN used as reference neuron to determine the triggering events, resulting in *NxN* PSF and PSTH models for all the available MNs. In addition, the same procedure was repeated for five surrogate MN pools per condition, in which the firing times of each MN were randomly shuffled while preserving the individual MN firing rate distributions.

PSF captures the instantaneous firing rate over time, offering a continuous representation of the temporal profile of synaptic input (Türker & Cheng, 1994; Powers & Türker, 2010). Excitatory input events were defined as increases in the instantaneous firing rate. The instantaneous firing rate modulation (pps) was estimated from the excitatory events in the PSF as the amplitude difference between the maximum and minimum CUSUM values in a window ranging from -5ms to 35ms (interval defined to include all potential increases in firing rate around the trigger) (**Figure 2A**). From the PSTH, synchronization events were estimated as the amplitude difference between the maximum and minimum CUSUM values in the same window (**Figure 2B**). To identify genuine excitatory and synchronization events separable from background noise, only the amplitude values higher than those obtained from the average of the surrogate MN pools were retained, following an adapted implementation of the error-box method in which the surrogate MN spike trains are used as the pre-stimulus window (Turker & Powers, 2005). The amplitude values for those MN pairs which were non-significant (i.e. excitation size smaller than the CUSUM error-box) were not considered. The synchronization values obtained from the PSTHs were normalized by dividing them by the total number of firings included in each PSTH. Excitatory or inhibitory modulations in the pre-trigger window were not analysed in this study, due to the potential contamination from the previous firing instants.

Matrices for the resultant instantaneous firing rate modulations (from PSF) and synchronization values (from PSTH) for all the combinations of MN pairs were generated (**Figure 2C, D**). The estimation of instantaneous firing rate modulation amplitude is unbiased by the MN firing rate and its distribution across the MN pool (Powers & Türker, 2010). Then, for each MN, the overall instantaneous firing rate modulation and synchronization amplitudes were estimated by averaging the instantaneous firing rate modulation locked to other MN firings (rows of the matrix), respectively. These estimates aim to answer the question: how does *the firing rate and synchronization of the target MNs change at the time instants when other MNs in the pool fire?*

Finally, for each subpool with MNs with the fastest and slowest firing rates, the instantaneous firing rate modulation and synchronization amplitudes were estimated as the average of the resultant from individual MNs within the subpool, an estimate which aims to answer the question: how does *the firing rate and synchronization of the target MN subpool change at the time instants when other MN subpools fire?*

#### Experimental data

Part of the dataset used in this study was originally described in a previous paper (Abbagnano *et al*., 2025). Fourteen healthy adults (28 ± 4.63 years; 12 males, 2 females) with no reported neurological or musculoskeletal impairments participated in this study.

Participants were seated comfortably with their dominant leg securely fastened to an ankle dynamometer using adjustable straps. They were asked to perform isometric dorsiflexion contractions of the Tibialis Anterior (TA) muscle with visual feedback at different force levels normalized to their maximum voluntary contraction (MVC). These contractions followed a trapezoidal force profile, consisting of a linear ramp-up phase (0% to target MVC at 5% MVC/s), a sustained plateau at the target level with a duration of 60 s, and a ramp-down phase returning to 0% MVC at the same rate. Four target force levels were tested: 5%, 10%, 20% and 30% MVC. A 2-minute rest interval was provided between trials to minimize fatigue. Testing across multiple contraction levels allowed us to examine whether the differential entrainment effects observed in simulations generalize across different MN pool configurations, since increasing force alters both the number of active MNs and their firing rates, thereby changing the relationship between MN firing properties and the frequencies of common input that would produce entrainment.

High-density surface electromyography (HDsEMG) signals were recorded from the TA muscle of the dominant leg using a 256-electrode grid (26 rows and 10 columns, 4 mm inter-electrode distance; OT Bioelettronica, Italy). EMG signals were acquired in monopolar configuration and amplified using a biosignal amplifier (Quattrocento, OT Bioelettronica, Torino, Italy). Data were sampled at 2048 Hz and band-pass filtered between 10 and 500 Hz. Simultaneously, force data were collected using a force transducer (TF-022, CCT Transducer s.a.s). These signals were digitized at 2048 Hz using the same amplifier.

The HDsEMG signals recorded during the experimental tasks were decomposed offline to extract the firing activity of individual motor units (MUs) using a convolutive blind source separation algorithm (Negro *et al*., 2016; Avrillon *et al*., 2024). To ensure data reliability, only spike trains with a silhouette (SIL) value of 0.90 or higher were retained (Holobar *et al*., 2014; Negro *et al*., 2016). Following automatic decomposition, all spike trains were manually inspected and edited to correct potential errors (Murks *et al*., 2025). All spike times were realigned to the onset of the motor unit action potential (MUAP) (Pascual-Valdunciel *et al*., 2025). The MN firing properties (iFR and 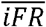) were computed for the spike trains corresponding to the plateau of the isometric contraction. Finally, MNs were grouped in quartiles based on their firing rate, following the same approach used in the simulation analysis (**Figure 4A**).

### Intramuscular coherence

Spectral methods, particularly coherence analysis, are widely used to assess stationary common input to MN pools (Dideriksen *et al*., 2018). Since in the experimental dataset we do not have direct access to the ground-truth common input projected to the MN pool, we computed intramuscular coherence (IMC) to estimate the common input received by the MN pool across frequency bands: theta [4, 8] Hz, alpha [8,12] Hz and beta [13, 35] Hz (Buzsáki, 2006; Negro & Farina, 2012*b*). The MN pool was randomly divided into two equally sized subgroups, and coherence between the resulting spike trains was calculated using MATLAB’s *Neurospec* 2.11 toolbox (Halliday, 2015). This procedure was repeated 50 times, each with a different random configuration of MUs. The final IMC estimate was obtained by averaging the coherence spectra across all iterations (Abbagnano *et al*., 2025).

### Statistical analysis

To test whether the MN-firing locked method captured the entrainment effects described above, we used Pearson’s correlation to compare, across common input frequencies (CIF_f_), the synchronization profile of each MN subpool with the instantaneous firing rate modulation profile of the complementary subpool.

For the experimental data, MN firing features 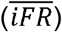 were compared across subjects using a two-way repeated measures ANOVA with MVC level and MN subpool as factors. The effects of these factors on instantaneous firing rate modulation were assessed using the same statistical approach. Post-hoc pairwise comparisons were conducted using paired t-tests with Bonferroni correction for multiple comparisons. Data normality and sphericity was assessed using the Shapiro-Wilk test and the Mauchly’s test.

For the relationship between instantaneous firing rate modulation amplitude and IMC, we used linear regression models. For each frequency band (theta, alpha and beta), we built a linear model with IMC amplitude as the dependent variable, instantaneous firing rate modulation amplitude as a continuous predictor, and contraction level (MVC) as a categorical predictor. Model goodness-of-fit was quantified using the coefficient of determination (R^2^). Statistical significance was set at α = 0.05 for all tests. All data processing and statistical analyses were performed using MATLAB (R2024b, The Mathworks Inc.) and Jamovi software.

## RESULTS

This section is divided into three parts. First, we used computational simulations to characterize the entrainment effects of common input frequency on different MN subpools. Then we demonstrate how this frequency-dependent entrainment effect translates into different firing rate dynamics analysed with the MN-firing locked method. Finally, we applied the MN firing locked analysis to experimental data from human TA muscle to investigate MN subpool sampling properties and their relationship with frequency-specific common inputs.

### Motoneuron simulations

#### Common input frequency entrains MN firings based on their firing properties

To characterize MN entrainment with common input, we first generated computational simulations of a MN pool receiving low-frequency common input that contributes to the steady MN firing observed during constant-force contractions, while an additional source of common input was projected to the MN pool at specific frequencies for different scenarios (**Figure 1A**). We systematically varied MN biophysical properties (from small to large S-type) as well as the frequency of the common synaptic input received by each MN in the pool. Instead of using a global synchronization measure of the full MN pool, our goal was to assess the synchronization of MN subpools with different firing rates. We selected the whole MN pool, and two MN subpools containing the active MNs with the fastest and slowest firing rates (first and last quartiles), and we computed the average MN-MN synchronization from PSTH normalized to the subpool number of firings.

The analysis revealed that the synchronization within MN subpools depends on the frequency of the common input received. MNs tended to fire synchronously when common input was delivered at frequencies closer to their firing rate or its harmonics (**Figure 3A, B**). The slowest MNs in the active pool firing approximately at 12.0±0.07 Hz increased synchronization when input was delivered between 9 Hz and 13 Hz, with a maximum at 12 Hz. Synchronization then decreased at higher frequencies until it increased again within the range of 21 to 29 Hz, corresponding to the first harmonic frequency band of average firing rate. In contrast, MNs with higher firing rates (16.22±0.3 Hz) maintained relatively low stable synchronization across frequencies, with a noticeable increase between 15 to 17 Hz and a maximum at 16 Hz, again aligned with their average firing rate. Synchronization increased in the frequency band from 31 to 34 Hz, the first harmonic band of the firing rate for this MN subpool. This effect is illustrated in **Figure 3B**, which shows how MN synchronization increases when the frequency of the common input approaches the firing rate of the MN pool or its multiple.

**Figure 3.**
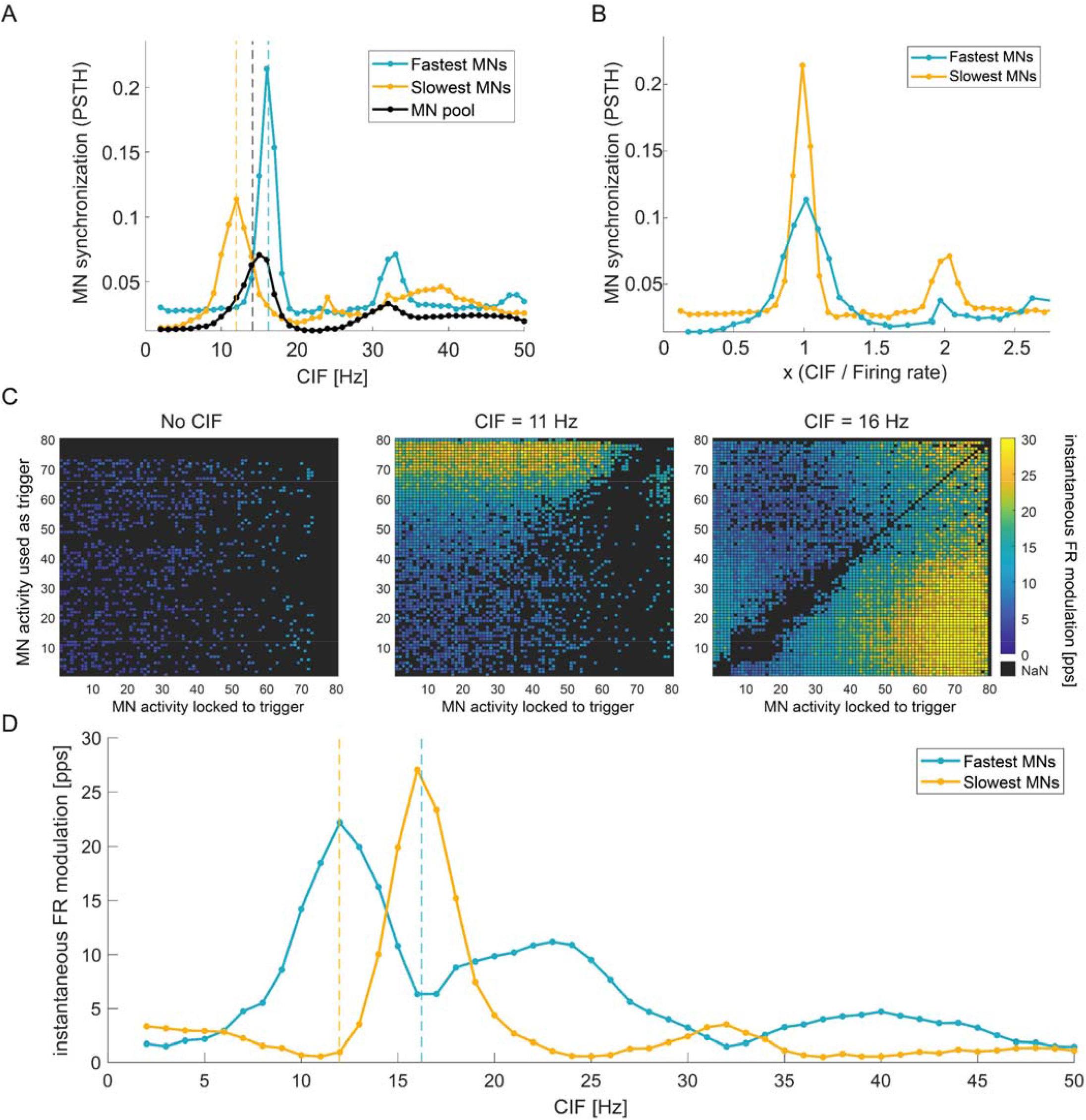
MN simulations analysis. **A**. MN synchronization (estimated from PSTH) across common input frequencies (CIF_f_) received by each MN sub-pool. Vertical dashed lines represent the mean firing rate of the colour-coded MN subpool. **B**. MN synchronization (estimated from PSTH) across common input frequencies (CIF_f_) received by each MN sub-pool. The x-axis (CIF_f_) was normalized to the mean firing rate for each MN subpool. **C**. Heatmaps of the instantaneous firing rate modulation amplitude across simulations with MN pools receiving common input at different frequencies. MNs are sorted by decreasing firing rate. **D**. Instantaneous firing rate modulation amplitude shown by the target MN sub-pool when the firings from the MN sub-pool based with the most different firing rate are used as trigger for the PSF.

When examining the MN synchronization across the entire MN pool, results showed a peak at 15 Hz, which matched the mean firing rate of the pool (14.15±0.2 Hz). Notably, the synchronization profile of the whole pool closely resembled that of the fastest MN subpool, while the entrainment of the slowest MNs was completely masked since no synchronization increase was detectable at frequencies matching the firing rate of the slowest subpool.

These findings support the hypothesis that common input frequency entrains MN firings based on their intrinsic firing properties. When the common input frequency, or one harmonic, approximates the firing rate of an active MN subpool, these MNs become entrained with the input. Instead, pooling the activity of groups of MNs with different firing rates masks the behavior of MN subpools.

#### High-frequency input induces different firing dynamics across MN subpools

We now exploited the entrainment effect to test whether the MN-firing locked method can capture the firing dynamics of MNs with different firing rates. Without any a priori knowledge of the common input frequency, we used the firing instants of entrained MNs as approximated temporal markers of common input, to apply a PSF analysis to pairs of active MNs from the previous simulations with varying common input frequencies (**Figure 2A, C**).

The MN-firing locked analysis revealed heterogeneous instantaneous firing rate modulations across MNs and simulations, depicted in the heatmaps in **Figure 3C**. The method captured transient increases in firing rate time-locked to the firing activity of other MNs, reflecting excitatory common input (**Figure 2A**). These modulations were not uniformly distributed across the pool. Instead, clusters of increased colour density appeared in specific regions of the heatmap for simulations where the common input frequency matched the firing rate of a specific MN subpool or its harmonic. For instance, at CIF_f_ = 11 Hz (**Figure 3C middle**), a cluster of increased colour intensity appeared in the top region of the heatmap: a wide range of MNs showed larger firing rate increases when the slowest MNs were used as time-locking events. Other input frequencies revealed different patterns. For example, (CIF_f_ = 16 Hz) revealed a cluster in the bottom right region, with increased colour intensity and density representing larger increases in firing rate for faster MNs in the active pool when the firing activity of slower MNs in the pool served as the time-locking trigger.

The MN-firing locked analysis yielded multiple data points from MN pairs that were difficult to interpret individually, thus, we reduced the dimensionality of the heatmaps by grouping into two different MN subpools comprising the MNs with the fastest and slowest firing rates. Specifically, we analysed the instantaneous firing rate modulation amplitude time locked to the firing activity of the complementary MN subpool. For the fastest MNs, we reported their transient modulation time locked to the slowest MNs’ firing activity, while for the slowest MNs, we reported their transient firing rate modulation time locked to the fastest MNs’ firing activity (**Figure 3D)**. The MN-firing locked analysis revealed different instantaneous firing rate modulation across MN subpools and common input frequencies. The slowest MNs in the pool (12.0±0.07 Hz mean firing rate) showed transient increases in their firing rate time-locked to the firing activity of faster MNs pool (16.2±0.3Hz mean firing rate) when the common input frequencies were delivered in specific frequency bands ([7,14] Hz and [19, 30] Hz). Conversely, the fastest MNs exhibited larger increases in their firing rate time-locked to the firing activity of the slowest MNs for the input frequency bands below 7Hz and within the [15,18] Hz band.

These firing rate modulation patterns resemble the MN entrainment properties reported in the previous section. To quantify this relationship, we correlated the synchronization and firing rate modulation profiles across common input frequencies. For each MN subpool, the entrainment profile predicted the firing rate modulation observed in the complementary subpool: the synchronization profile of the slowest MNs correlated with the firing rate modulation of the fastest MNs (r = 0.72, p < 0.001), and the synchronization profile of the fastest MNs correlated with the firing rate modulation of the slowest MNs (r = 0.89, p < 0.001). Altogether these findings suggest that the firing activity of the MN subpool, when used as triggering event in the MN-firing locked analysis, can be used to infer the common input modulatory effects on the complementary MNs within the pool.

### Experimental dataset

Individual MN activity from humans can be inferred through electrophysiological recordings such as HDsEMG. However, in these cases, the ground-truth common input is not known. The MN-firing locked method presented in this work could provide a window towards understanding the relationship between the individual MN activity and the frequency of the common input in experimental conditions. In this section, we applied the MN-firing locked method to MN pools activity recorded from the TA from healthy individuals using HDsEMG during voluntary contractions.

A total of 1557 MUs were decomposed and retained for analysis across all subjects and contraction levels. The average number of MNs per subject was 24.9±6.5 (5% MVC), 29.4±9.8 (10% MVC), 29.6±10.4 (20% MVC) and 27.3±10.6 (30% MVC). For each subject and contraction level, MNs were grouped by firing rate, and the two quartiles corresponding to the MNs with fastest and slowest firing rates in the active MN pool were selected for analysis (first and last quartiles). Under the onion skin principle, MNs with higher firing rates correspond to smaller, earlier recruited units, while those with lower firing rates represent larger and later recruited units (Piotrkiewicz & Türker, 2017) (**Figure 4A**). For each contraction level, MN firing rates of the two subpools selected differed significantly across subjects (RM ANOVA; the fastest MNs vs the slowest MNs): 5% MVC (11.6 vs 9.0 pps; p < 0.001), 10% MVC (12.7 vs 9.8 pps; p < 0.001), 20% MVC (13.9 vs 10.5 pps; p < 0.001) and 30% MVC (15.1 vs 11.0 pps; p < 0.001).

**Figure 4.**
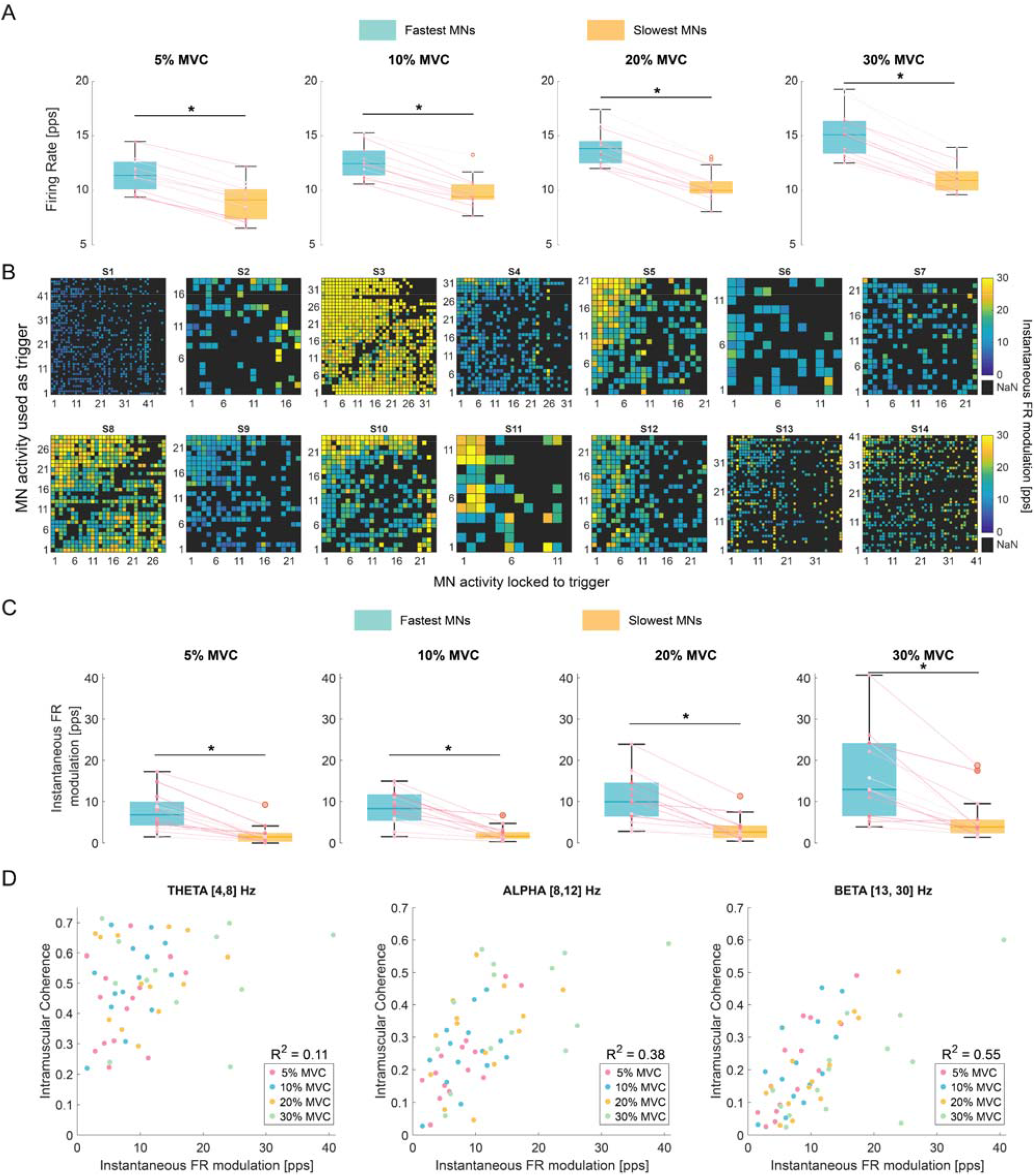
Experimental results. **A**. Firing rate of the two MN-subpools grouped across subjects and contraction levels. Coloured dots represent individual subject data. **B**. Instantaneous firing rate modulation heatmaps for all subjects during 30% MVC dorsiflexion. MNs are sorted by decreasing firing rate. **C**. Instantaneous firing rate modulation time-locked to the opposite size MN group firing activity. **D**. Intramuscular coherence (IMC) and instantaneous firing rate modulation amplitude for the fastest MNs in the active pool locked to the firing activity of the slowest MNs. Each contraction level (MVC) is colour coded. R^2^ represents the linear regression model goodness-of-fit. * Denotes statistical significance (p<0.05).

#### MN-firing locked analysis reveals different modulation of firing rate across MN subpools

The MN-firing locked analysis revealed the presence of heterogeneous instantaneous firing rate modulations across MNs and participants (**Figure 4B)**. The PSF-CUSUM analysis revealed that, on average, 35±15 % of MNs transiently increased their firing rate when their spiking activity was time-locked to the firing activity of other MNs. This proportion is higher than that observed in the MN simulations (17%) in which no high-frequency common input was delivered (CIF_0_). This result suggests that the transient modulations of MN’s firing rate in the TA are elicited by high-frequency common inputs.

Clusters of increased colour intensity, reflecting MNs receiving excitatory inputs that translate into instantaneous firing rate increases, were visible in specific regions of the heatmaps for most subjects (**Figure 4B**, e.g., S5, S8, S10). This pattern was particularly evident in subject S5, who showed a distinct cluster in the top left region where faster MNs (left columns) exhibited larger firing rate increases when time locked to the firing activity of slower MNs. In contrast, other subjects such as S1 and S2 did not reveal distinct visual clustering patterns, with instantaneous firing rate modulation remaining relatively absent across MN pairs. This inter-subject variability likely reflects differences in the frequency content and amplitude of common synaptic inputs across individuals.

Analysis of MN subpools revealed an asymmetric pattern. When the firing activity of the slowest MNs was used as triggering events, the fastest MNs exhibited larger increases in firing rate compared to the complementary condition. Contrarily, when the fastest MNs were used as triggers, the slowest MNs showed smaller firing rate increases (RM ANOVA; p<0.001, **Figure 4C)**. This group-level pattern was consistent across all contraction levels tested, with differences observed in the instantaneous firing rate modulation amplitude across MN subpools (RM ANOVA; the fastest MNs vs the slowest MNs): 5% MVC (7.6±4.6 vs 2.0±2.4 pps; p < 0.001), 10% MVC (8.4 ± 4.0 vs 2.2±1.8 pps, p < 0.001), 20% MVC (10.7±6.0 vs 3.4±2.9 pps, p = 0.001) and 30% MVC (16.0±1.3 vs 5.9±5.5 pps, p = 0.006). Furthermore, contraction level had a significant effect on the instantaneous firing rate modulation amplitude (RM ANOVA, p<0.001).

#### High frequency common input drives transient firing rate dynamics

In the simulations, the common input delivered to the MN pool was known and controlled, allowing direct attribution of the observed firing rate modulations to specific input frequencies. In the experimental dataset, however, the common input is not known *a priori*. To determine whether the experimental firing rate modulations reflected frequency-specific common input, as predicted by the simulations, we estimated the common input to the MN pool using intramuscular coherence (IMC).

To investigate the interaction between instantaneous firing rate modulation and the common input frequency, we built a linear regression model to predict the IMC. For different frequency bands we used the instantaneous firing rate modulation amplitude from the fastest MNs in the pool as a continuous predictor and the contraction level (MVC) as a categorical independent variable. (**Figure 4D**). The models revealed a moderate interaction between IMC amplitude and instantaneous firing rate modulation for the alpha (R^2^ = 0.38, p<0.001) and beta bands (R^2^ = 0.55, p<0.001), while no linear relationship was present for the theta band (R^2^ = 0.11, p =0.35). Overall, these findings suggest a relationship between the instantaneous firing rate modulation shown by the fastest active MNs and the strength of common input for frequencies above 8 Hz. Together, these results support the feasibility of the MN-firing locked method in effectively capturing the transformation of high-frequency common input, particularly in the alpha and beta bands, into transient firing rate dynamics across different MN subpools.

## DISCUSSION

Through computational simulations and experimental data, we showed that MNs with different firing properties in the active pool transform differently the high-frequency common input into temporal firing rate dynamics. Our results also highlight the importance of considering the distribution of MN properties in the pool, particularly firing rate, when interpreting population-level metrics such as global MN synchronization. Population-level analyses often treat the MN pool as integrating the shared synaptic drive in a broadly similar way across neurons, so that the overall output is approximately linear. This linearization is likely optimal for robust force transmission, where the aggregate pool output is what matters functionally. However, for common input at frequencies above the force bandwidth, this pooling masks how individual MN subpools sample the input differently depending on their firing rates, as demonstrated in this study. While these hidden dynamics may have limited functional consequences in healthy steady contractions, there are important scenarios, such as motor disorders, where the entrainment of specific MN subpools with pathological oscillatory inputs may become critical. Additionally, the different vulnerability of MN types that is characteristic of some neurological disease conditions, such as motor neuron disease (MND), demands more robust and sensitive analysis at the subpool level. The methodology and results presented here open new avenues for studying MN physiology in both healthy and pathological conditions.

### Motoneuron firing entrainment driven by common input frequency

Computational simulations revealed that MNs tend to fire in synchrony with the oscillations of the input frequency matching their firing rate, consistent with previous studies (Lowery & Erim, 2005). This global behavior is expected when common input is uniformly projected to a uniform MN pool. However, our study extends beyond previous work by examining how individual MN subpopulations within the pool respond to frequency specific inputs, rather than focusing solely on the pooled activity of all the MNs. A key finding is that MN subpools with different intrinsic properties exhibited different entrainment to specific frequencies of common input. Importantly, this entrainment phenomenon occurred not only at frequencies matching MN firing rates but also at harmonic frequencies. This reflects the undersampling property of individual MNs, a characteristic non-linearity whereby MNs cannot follow input frequencies exceeding their firing rate under physiological conditions (Lowery & Erim, 2005; Ibáñez *et al*., 2021). However, their activity can become entrained to sub-periodic components of faster oscillatory inputs, effectively locking to harmonics or subharmonics of the driving frequency.

From this study alone it is not possible to fully discern the functional role of the differential entrainment across MN subpools. The common input frequencies that produce the strongest subpool-specific entrainment (alpha and beta bands) have limited direct effects on steady-state force modulation (Zicher *et al*., 2023). Thus, the heterogeneous synchronization patterns reported here across individuals may represent an unavoidable consequence of how individual MNs with different firing rates process oscillatory common input, rather than a mechanism actively exploited by the nervous system for motor control. Alternatively, these frequency bands are associated with a range of sensorimotor functions beyond force control, including corticospinal communication, sensorimotor integration and responsiveness to perturbations (Baker, 2007). During sustained contractions, descending commands, spinal circuits and peripheral inputs contribute to oscillatory activity that may serve functions not yet fully characterized. The heterogeneous transformation of these inputs across MN subpools, as revealed by our method, may therefore reflect functionally meaningful differences in how MN subpopulations process relevant signals, rather than simply an epiphenomenon of MN properties.

Differential entrainment may become functionally significant when pathological oscillatory inputs arrive at frequencies closer to the force bandwidth. In pathological tremor, oscillatory common input from 4 to 8 Hz overlaps with or falls near the subharmonics of MN firing rates in the active pool (Gallego *et al*., 2015). Our findings predict that MN subpools with firing rates matching the tremor frequency or its harmonics would become preferentially entrained. Similarly, in neurodegenerative conditions, such as MND, where fast-type MN denervation and compensatory reinnervation by surviving slow-type MNs reshapes both MN pool composition and the synaptic drive delivered to muscles (Huh *et al*., 2021), the entrainment landscape could shift in multiple ways, even years before detectable gross functional motor impairment. The MN-firing locked method presented here provides the analytical framework to test these predictions, extending common input characterization to reveal how individual MN subpools are affected in pathological conditions.

Although single- and two-compartment computational models have been shown to successfully replicate changes in MN firing under constant common input (Cisi & Kohn, 2008; Powers & Heckman, 2017), we acknowledge that our simulation results are based on simplified models incorporating common input without neuromodulatory influences or feedback circuits that are present *in vivo*, such as persistent inward currents, recurrent inhibition, or Ia reciprocal excitation among others (Williams & Baker, 2009; Yavuz *et al*., 2015; Mesquita *et al*., 2024). Future modelling efforts should aim to incorporate these additional physiological mechanisms for further interpretation of experimental data.

### Instantaneous firing rate modulation across MN subpools

We present an application of PSF-based analysis to large MN active pools decomposed from HDsEMG, demonstrating that this method is sensitive to detect subtle differences in excitatory inputs through transient firing rate modulations across individual MNs. Analysis of the experimental dataset revealed consistent asymmetry in instantaneous firing rate modulation across MN subpools. Smaller MNs, characterized by higher firing rates, exhibited significantly greater increases in firing rate when their activity was time-locked to the firing of slower firing MNs. This pattern was observed across all tested contraction levels and proved robust to inter-subject variability at the group level, suggesting a fundamental organizational principle of MN pool dynamics.

Traditionally, a large portion of firing rate variability during constant force tasks has been attributed primarily to the interaction between intrinsic MN properties and independent synaptic noise, with faster firing MNs exhibiting lower coefficients of variation (Moritz *et al*., 2005). However, our results demonstrate that within this variability, there is systematic, non-random modulation reflecting the integration of common synaptic input. These structured firing rate dynamics reveal that MN firing rate variability during sustained contractions —commonly quantified with the coefficient of variation— contains information about frequency-specific common inputs beyond simple noise or independent fluctuations.

Force level significantly influenced the magnitude of firing rate modulation. Increasing contraction levels produced larger instantaneous increases in firing rate time-locked to the firing of other MNs. This force dependence likely reflects the concomitant increase in amplitude for both low frequency input, contributing to force production, and high frequency input, which together produce stronger modulations of MN excitability. The larger number of MNs recruited at increasing forces could influence these results, however, the MN count included in the analysis did not consistently increase across the contraction levels.

The findings reported in this study remain consistent with the concept of linear transmission of a common input to the MN pool (Farina *et al*., 2014). A uniformly distributed input across the entire MN pool can still result in varied sampling characteristics among MN subpools. Nevertheless, when considering the collective activity of the entire MN pool, the input is transmitted in a linear fashion, despite the non-linear dynamics observed at the level of individual MNs or subpools (Farina & Negro, 2015).

### High frequency common input drives transient firing rate dynamics

The MN-firing locked and PSF methods do not imply any causality between the two events sampled defined as the firing activity of the MN used as trigger and the firing rate modulation of the tested MN. A possible interpretation of the observed asymmetric distribution of firing rate modulations in experimental data is the presence of recurrent excitatory circuits projecting from larger to smaller MNs in the active pool. However, this interpretation is inconsistent with known spinal microcircuits in mammals. While recurrent excitation has been proposed as a mechanism for MN-MN interactions, in vitro studies suggest that such excitation is preferentially directed among fast-type MNs, not from fast to slow or slow to slow types (Bhumbra & Beato, 2018; Özyurt *et al*., 2024).

An alternative hypothesis to direct causality considers the common input that drives the firing activity of the MNs as the source responsible for the findings reported with experimental data. When the frequency of the common input aligns with the firing rates of the slowest MNs active in the pool, these MNs become entrained and serve as effective samplers of the shared synaptic drive, as suggested in previous studies (Lowery & Erim, 2005). Consequently, using their firing activity as temporal triggers in the MN firing-locked analysis reveals the excitatory influence of the common input on other MNs within the pool.

Results from the application of the MN-firing locked analysis to MN simulations are aligned with this hypothesis. Importantly, these findings were observed despite the models not including any connectivity between individual MNs. The instantaneous firing rate modulation at specific frequencies correlated with the entrainment of the MN subpool firing activity used as reference. Particularly, only inputs delivered at physiologically relevant frequencies, specifically within the alpha and beta bands, reproduced the experimental patterns of instantaneous firing rate modulation where the fastest MNs exhibited larger modulations compared to the slowest MNs in the pool (Baker, 2007; Cabral *et al*., 2024). Both frequency bands are well established components of descending and spinal common drive to the MN pool during voluntary contractions (Farmer *et al*., 1993; Graef *et al*., 2025). Further analysis of common input from experimental recordings showed a linear relationship between instantaneous firing rate modulation and intramuscular coherence in the alpha and beta bands. This convergence of computational simulations and experimental observations indicates that high frequency common input is likely responsible for the transient firing rate modulations captured by the MN-firing locked method.

A key aspect of the experimental findings is the variability observed among individuals. While some individuals showed pronounced differences in the instantaneous firing rate modulation across different MN subpools, others did not exhibit this asymmetric distribution. Interestingly, these differences were consistent with the distribution of common input at high frequencies estimated through IMC. This inter-subject variability would reflect the nervous system’s flexibility in achieving fine motor control by employing diverse neural strategies across individuals (Pascual-Valdunciel *et al*., 2025; Kim, 2025; Sahinis *et al*., 2025). It suggests that the human nervous system can modulate the frequency content of inputs to the MN pool in different ways, allowing for different configurations across individuals while still maintaining consistent functional outcomes.

## Additional Information

### Data Availability statement

Data from this study will be made available to qualified investigators upon reasonable request to the corresponding author. The computational model and analysis code used in this study were implemented in Julia. The code is available upon request from the corresponding author.

### Author contributions

All authors approved the final version of the manuscript. All authors agree to be accountable for all aspects of the work. All authors qualify for authorship, and all those who qualify for authorship are listed

A.P-V.: Conceptualization, Data acquisition, Investigation, Analysis, Visualization, Writing, Funding Acquisition.

J.Y-M.: Conceptualization, Investigation, Analysis, Visualization, Writing.

E.A.: Data acquisition, Analysis, Writing.

N.T.C.: Software, Analysis, Writing.

F.N.: Investigation, Analysis, Visualization, Writing.

M.G.O.: Investigation, Analysis, Visualization, Writing.

D.F.: Conceptualization, Investigation, Analysis, Visualization, Writing, Resources, Funding Acquisition.

J.I.: Conceptualization, Investigation, Analysis, Visualization, Writing, Resources, Funding Acquisition.

